# Generative Adversarial Networks Conditioned on Brain Activity Reconstruct Seen Images

**DOI:** 10.1101/304774

**Authors:** Ghislain St-Yves, Thomas Naselaris

## Abstract

We consider the inference problem of reconstructing a visual stimulus from brain activity measurements (e.g. fMRI) that encode this stimulus. Recovering a complete image is complicated by the fact that neural representations are noisy, high-dimensional, and contain incomplete information about image details. Thus, reconstructions of complex images from brain activity require a strong prior. Here we propose to train generative adversarial networks (GANs) to learn a generative model of images that is conditioned on measurements of brain activity. We consider two challenges of this approach: First, given that GANs require far more data to train than is typically collected in an fMRI experiment, how do we obtain enough samples to train a GAN that is conditioned on brain activity? Secondly, how do we ensure that our generated samples are robust against noise present in fMRI data? Our strategy to surmount both of these problems centers around the creation of surrogate brain activity samples that are generated by an encoding model. We find that the generative model thus trained generalizes to real fRMI data measured during perception of images and is able to reconstruct the basic outline of the stimuli.

## I. INTRODUCTION

The goal of neural decoding is to characterize some distribution *p*(*X*|*V*), where *X* is a certain class of percepts (e.g. a class of objects, a direction, an image, etc.) and *V* is a set of neural observables measured under some procedure (e.g. fMRI voxel measurements). In the context of vision, we might be interested in some property of an image or the image itself. When *X* is an image, *p*(*X*|*V*) would be a useful interpretive tool by allowing one to sample pictures associated with a measured brain state. Generating such samples— a problem known as image reconstruction—is difficult for several reasons. First, even measurements of brain activity that permit perfect “identification” [1] of images may only partially characterize those images. Second, invariances to image detail are built into the brain’s visual representations from the earliest stages of visual processing. These invariances are necessary for object recognition, but induce a one-to-many relationship between brain states and images. Meaningful reconstruction thus requires prior information that can be combined with information from the brain to constrain the decoded images. Previous methods for image reconstruction [2], [3] have utilized massive collections of photographs as an implicit prior. Under this approach only images that have previously been photographed can be decoded, and it is often not clear what parts of the decoded image are strongly or poorly constrained by the neural representations. These can be unwelcome limitations when considering reconstruction of mental images, for example, since mental images may potentially depict things that have never been seen before.

Here, we take several steps toward replacing large databases of images with a trainable and sufficiently generic generative model. Generative models characterize a probability distribution *p*(*X*), either explicitly through parametrization of a known distribution or implicitly by providing a sampling process with which to obtain samples from *p*(*X*). A class of generative models known as generative adversarial networks (GANs) have received a lot of interest lately for their ability to produce convincing-looking image samples [4]–[6]. These generative networks have several desirable properties. First, they do not require strong assumptions regarding the form of *p*(*X*) because it is only defined implicitly by the model. GANs implicitly represent *p*(*X*) by first sampling from an arbitrary distribution *p*(*Z*) and then transforming it through a nonlinear function (represented by a deep generator network) *G_X_* to make it look like *p*_data_(*X*) to a discriminator function *D_X_* trained to distinguish *p*_model_(*X*) from the real image distribution *p*_data_(*X*). Secondly, the discriminator, being itself a deep network jointly trained with the generator network, is in effect a trainable objective function. Therefore, the discriminator is left to discover what (statistical) measures best characterize the data.

Importantly, GANs have been generalized to produce image samples from a conditional distribution *p*(*X*|*Y*) where *Y* is typically a low-dimensional noise-free code such as an object category embedding. However, it has also been suggested (and demonstrated in content transfer applications) that the conditional GAN framework could be extended to conditions of multiple modalities [6]. In this study we condition a GAN on high-dimensional, distributed brain-like codes *C* and examine the prospects of such a conditional generative model for stimulus reconstruction. In principle, assuming that the code accurately characterizes specific image details, a conditional GAN should be able to learn to exploit this information to narrow the distribution *p*(*X*|*C*) around the target image. If successful, samples from *p*(*X*|*C*) would be faithful reconstructions of the target image.

We use *in silico* brain-like activity to provide a proof-of-principle demonstration that a conditional GAN can learn to use stimulus features encoded in a conditioning vector *C* to produce faithful reconstructions of a target image. We show that the variability of these reconstructions—which are ostensibly samples from *p*(*X*|*C*)—is directly related to the content and quality of the conditioning code. We then apply our approach to reconstruct target images from fMRI measurements of brain activity in human visual cortex. Applying our approach to real brain activity measurements is challenging due to the generally low signal-to-noise associated with these measurements, and because it is currently not feasible to obtain enough samples of brain activity to successfully train a conditional GAN. We overcome these challenges by using a voxelwise encoding model of activity in the visual cortex. The encoding model generates predictions of brain activity in response to any arbitrary stimulus. As we will demonstrate, we can use these predicted activity patterns to learn to denoise and compress raw brain activity measurements into a conditioning code *C*. We can further leverage the encoding models to generate tens of thousands of surrogate data samples that can be used to train a conditional GAN. We show that this approach then generalizes to reconstructions of natural scenes from human brain activity.

## Methods

Our method consists of three distinct components shown in Fig. 1. First, we build an encoding model. A deterministic encoding model makes some neural predictions 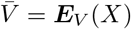 (whose elements may correspond to neurons, voxels, etc), where *E_V_* (*X*) is some mapping from stimulus space to neural space. Second, we reduce the dimensionality of the voxel activity vector *V* by utilizing the internal representations *C* = *E_C_*(*V*) of a denoising auto-encoder. Finally, we build a generative network conditioned on the condition vector *C*. We used the energy-based formulation of GANs [7] since it provided good sample quality and more stable training for all conditions. These three steps are detailed below.

**Fig. 1.**
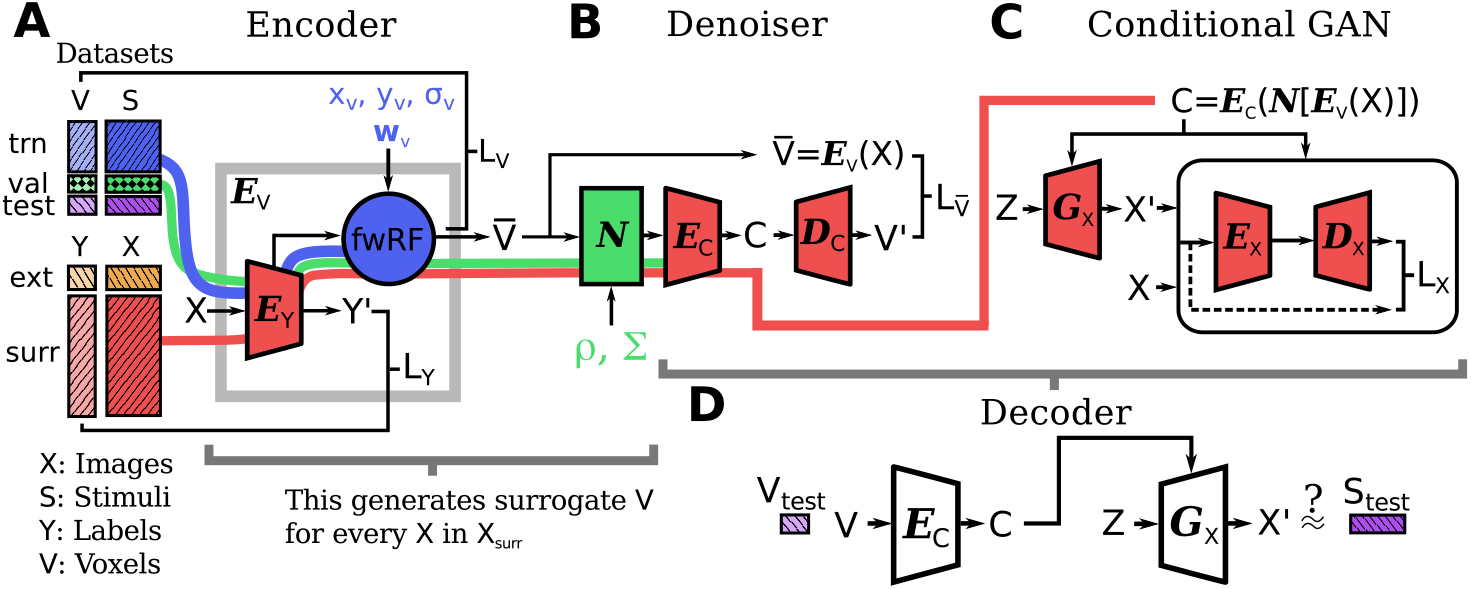
Summary of the model. The proposed strategy consists of three independently trained components: (A) A voxel-wise encoding model based on the fwRF, (B) a voxel denoising autoencoder and (C) a conditional generative network (here an ebGAN). (D) shows the components of the system used to sample images from *p*_model_(*X*|*C*=*E_C_*(*V*_test_)) at test time. *E, D* and *G* represent encoder, decoder and generator deep neural networks. *S* and *V* are stimuli and fMRI brain activity measures, respectively, from the vim-1 dataset. *X* and *Y* are images and categorical labels, respectively, from a separate image collection (cifar-10 or MNIST). Arguments to the loss functions *L* are indicated by the black brackets. The color of the various model components is matched to the data subset (labeled trn, val, test and surr) used to train their parameters; colored pipelines show the flow of data through the system during training (A-C). Models in (A) are trained sequentially (not jointly) from left to right i.e. first *E_Y_*, then the fwRF followed by a voxel noise model *N*. Then all the parameters in (B) are trained jointly followed by all parameters in (C). The surrogate voxel data used to train (B) and (C) is obtained by applying the encoding model (and noise model) to the input images of *X*_surr_.

### A. Encoding

In previous work, we have detailed an approach to encoding models called the feature-weighted receptive field (fwRF) model [8]. This approach makes several assumptions— foremost is space-feature separability—which heavily regularizes the training of the model. For our purpose here, it also has the additional benefit of introducing several easily interpretable and controllable parameters.

The fwRF model *E_V_* (*X*) (Figure 1A) can be separated into two components: a feature extractor *E_Y_* and a linear (or weakly nonlinear) feature-to-voxel regression (the blue circle labeled fwRF in Figure 1A). For the sake of demonstration, we used a very low spatial resolution deep neural network (DNN) as a feature extractor. This DNN consisted of five convolutional layers (with rectifier nonlinearity) and one fully-connected layer trained to recognize the 10 image categories of the cifar-10 dataset [9]. This resulted in a total of 2314 feature maps of various resolutions, which were all included in the fwRF model. The fwRF model applies to these feature maps a set of feature weights w*_v_* and a spatial receptive field (RF) parameterized by a size *σ_v_* and center location (*x_v_, y_v_*), where *v* indexes a particular voxel. The feature weights determine the importance of each feature map in the DNN to the predicted response of the voxel being modeled, whereas the spatial RF determines the region of the visual field where an individual voxel is most sensitive to the features with nonzero weights. Feature weights are learned via gradient descent, while spatial RF are learned via grid search over a set of candidate RFs. In this work, the search grid included 8 log-spaced sizes *σ_v_* between 0.7 and 6.0 degrees spaced 1.33 degrees apart (regardless of size) for a total of 1800 candidate spatial RFs. A compressive element-wise nonlinearity 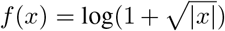 was applied after pooling over the feature maps, which resulted in slightly better prediction accuracy. The model for each voxel was run for 40 epochs of stochastic gradient descent with batch size of 200 and step size of 10^−4^ starting from an initial state of w*_v_* = 0.

### B. Denoising

Image features are encoded noisily and redundantly across populations of voxels in fMRI measurements of brain activity. This suggests that high-dimensional, noisy fMRI brain activity patterns can be compressed into a much lower-dimensional embedding. We used a simple denoising autoencoder to produce a highly informative and noise-resistant embedding of the brain activity. An autoencoder is a mapping *X′* = *D_C_*(*E_C_*(*X*)) where *E_C_*(*X*) provides some embedding of *X* into a (usually smaller or sparser) space *C*. The training objective involves minimizing the distance between *X′* and *X*. A denoising autoencoder uses the same objective but adds some noise to the input of the encoder *X′* = *D_C_*(*E_C_*(*N*[*X*])), where *N*[·] is some corruption process, which further regularizes the encoding and decoding procedure and increases robustness to corrupted inputs [10]. The target is the uncorrupted input *X*, and the goal is usually to discover some useful feature representations of the data.

In this work, we used a denoising autoencoder to denoise brain activity measurements *V* under the assumption that the outputs of the voxel encoding model 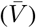 provide a suitable denoised target. Therefore, our autoencoder objective is

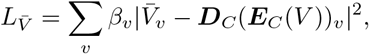

where *v* indexes the various voxels of *V* 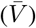 and *β_v_* is some relative importance factor for voxel *v*. In order to obtain enough data samples to effectively train the autoencoder, we used a surrogate data approach (note that this approach was also used to train the GAN, as will be seen below). First, we generated a large set of encoding model predictions 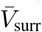 corresponding to images in a collection of natural scenes (here the cifar-10 dataset). These predicted brain activity vectors served as target activity patterns for training the denoiser. To create the surrogate brain activity patterns for training, we corrupted the prediction 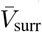 under a noise model

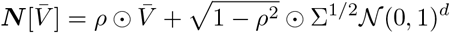

that is consistent with the validation accuracy (i.e., the correlation between the predicted and the experimentally measured brain activities for each voxel) and covariance structure (i.e., the correlation of residuals across the voxel population) of the real brain activity. Here, *ρ* = *ρ*_val_ and Σ^1/2^ are noise parameters estimated from a special validation set of brain activity measures, *V*_val_. Briefly, these noise parameters satisfy 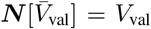. The training objective above was minimized by applying stochastic gradient descent to 59500 surrogate brain activity measurements. This process generalized extremely well to denoising *V*_val_.

### C. Decoding

To produce reconstructions of images from measurements of brain activity, we trained a GAN to generate samples from an implicit distribution *p*(*X*|*C*), where *C* is a conditioning vector derived from brain activity measurements obtained while a human subject observed a target image. The GAN training strategy does not minimize a single objective function, but iteratively updates a generator and discriminator network to minimize separate, opposing, objectives [4]. This process admits a unique global optimal solution when the generator produces samples from the (conditional) data distribution, i.e. *p*_model_(·) = *p*_data_(·), limitations due to realization notwithstanding.

For this work, we trained a conditional energy-based GAN (ebGAN) [7], which is a modification of the structure of a GAN that replaces the binary discriminator by a deep autoencoder network (note that the autoencoder used to train the ebGAN is completely distinct from the autoencoder we used to obtain conditioning codes for the ebGAN). In this formulation, the conditional generator *G_X_* (*Z*; *C*) and conditional autoencoder *D_X_*(*E_X_*(*X*; *C*); *C*) parameters are trained under their respective objectives

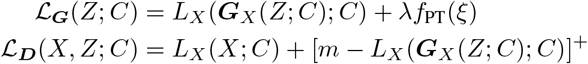

where 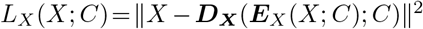 is a standard L2-norm loss for a deep autoencoder and [·]^+^ indicates a rectifier nonlinearity. Like in Ref. [7], the addition of a “pull-away” regularization term ƒ_PT_(ξ) to the autoencoder internal representation ξ greatly reduced the preponderance of mode collapse in our tests. *m* and λ were set to 1.0 and 0.1 respectively. Conditioning is realized by concatenation of the condition vector *C* with *every* feature map (replicated over spatial extent) of the generator and discriminator networks [6]. Here, *C* is a 128-dimensional vector and *Z* is sampled from a 920-dimensional normalized and uncorrelated Gaussian which is sliced and concatenated at various layers of the generator network. The training of the conditional generative adversarial network then followed standard practice.

Note that nowhere in the GAN training procedure do we specify explicitly that the generated images have to be close to the encoded images. However, by learning to fool the discriminator by matching the statistical features of images from the dataset, the generator learns to use stimulus information encoded in the conditioning vector *C* to its advantage. This results in sample images that are heavily constrained toward the image encoded in the brain activity measurements used to produce the conditioning vector. On the other hand, if a great amount of noise remains in the conditioning vector, the implicit distribution *p*(*X*|*C*) is weakly constrained by *C*, so that samples from this distribution would resemble those of the unconditional GAN. Therefore, the training procedure effectively decides whether or not to leverage the conditioning vector. One consequence of this is that a complete retraining of the GAN has to be performed for any change in the noise distribution or in the features encoded in the (synthetic) voxels.

All networks were implemented in theano [11] (lasagne [12]). To train the generator we used the same surrogate data used to train the denoiser. At this point, it is straightforward to generate an associated noisy condition vector *C* = *E_C_*(*N*[*E_v_*(*X*)]) for any image *X* in a large dataset. Using the previous 59500 surrogate data samples, we trained the generator and discriminator with the Adam optimizer for 400 epochs, with learning rates of 2.5 × 10^−4^ and 5.0 × 10^−4^ for the generator and discriminator respectively, alternating training between each batch of size 500. After 250 epochs, the learning rates were reduced by 1% each epoch until the end of training. The generated images from the training set usually started to be recognizable after over a hundred epochs. At test time, we constructed a conditioning vector by submitting a real brain activity measurement *V*_test_ to the trained encoder *E_C_*. This was then passed to the generator *G_X_* to generate image samples (see Fig. 1D). Such image samples are shown in Figs. 2A, 2E, 4 and 5). In this way, multiple reconstructions are obtained for each brain activity measurement.

**Fig. 2.**
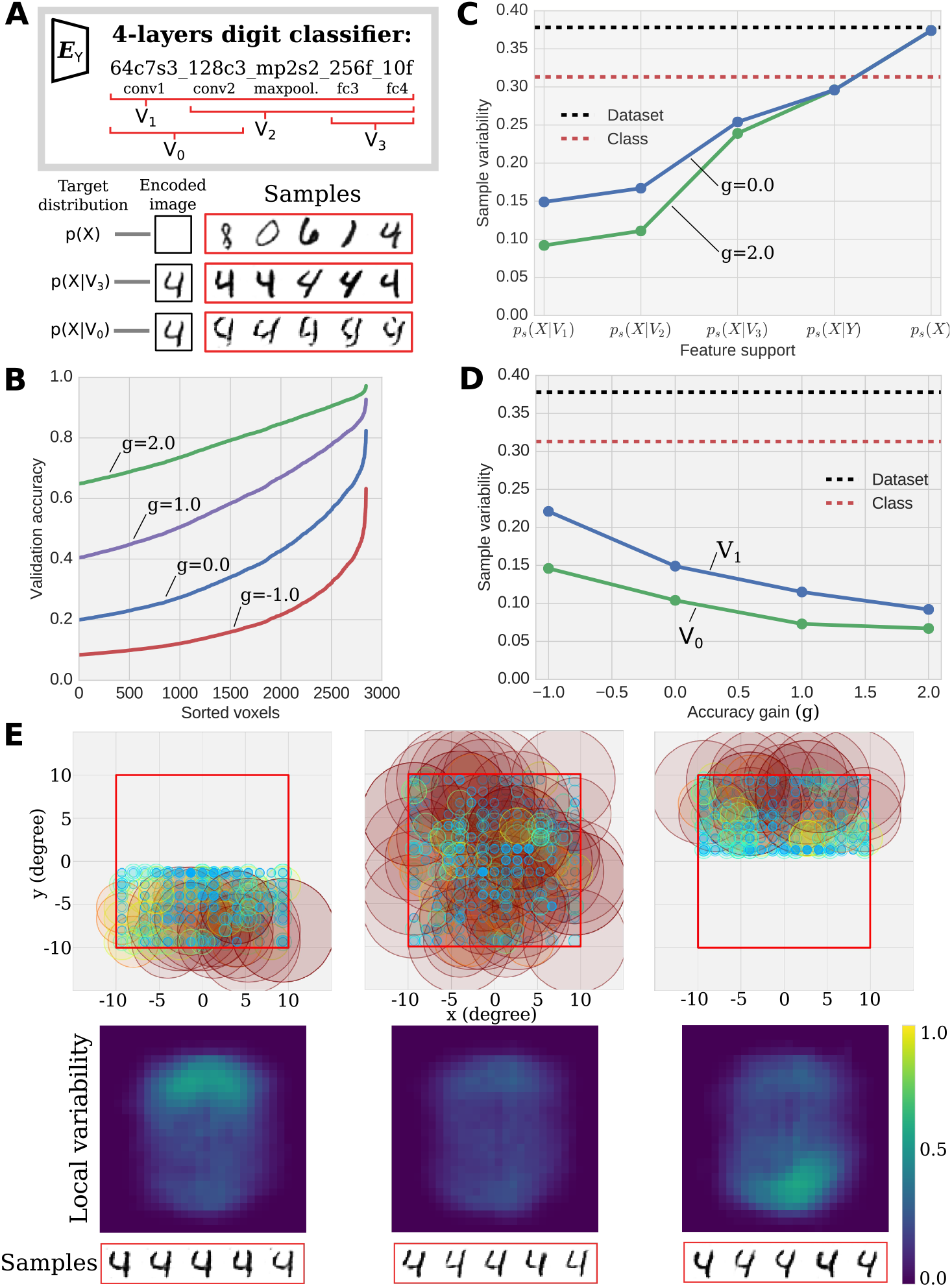
Control studies on a synthetic dataset. (A) We generated synthetic brain activity populations tuned to different combinations of layers in a DNN, *E_Y_*, that was trained to discriminate digits in the MNIST image collection. Synthetic activity patterns were generated using 4 distinct sets of features maps: *V*_0_ used only the first 2 convolutional layers; *V_i_* used all feature maps above and including layer *i*. The set of feature maps used to produce synthetic voxel responses is referred to as the population’s “feature support”. (Below) An example of generated samples based on the encoded stimuli. Samples from *p*(*X*) are unconditioned. The samples from *p*(*X*|*V*_3_) are conditioned on high-level features only and show a consistent “4” (a “4” was indeed encoded) with large variability in style. Samples from *p*(*X*|*V*_1_) for the same encoded image show much less variability (as shown at the very bottom of the figure). (B) Distribution of synthetic voxels model accuracy (i.e. how accurately the known encoding model for each synthetic voxel predicts its signal) with varying levels of noise. Each distribution is parameterized by a “gain” coefficient *g*. A positive (negative) gain means that the synthetic population includes voxels with more (less) accurate encoding models than the reference distribution (null gain). (C) *Effect of the encoding model feature support*. Sample variability is a measure of the variability of image reconstructions sampled from the implicit image distribution learned by the GAN. The two right-most points show the sample variability of the unconditioned *p*(*X*) and class-conditioned generator *p*(*X*|*Y*) (*Y* is a one-hot class label) relative to the unconditioned (black dashed line) and average class-wise variability (red dashed line) of images in the MNIST dataset. (D) *Effect of the model accuracy*. One can test the effect of model accuracy by choosing different distributions of voxel noise, as shown in B. As would be expected, synthetic voxel populations with little noise result in lower sample variability. (E) *Effect of the receptive field support*. Circles show the size and locations of RFs in three separate synthetic voxel populations. Populations with RFs restricted to a single visual hemifield lead to increased sample variance in the un-represented hemifield, while preserving the category of the represented digit. Results for each distinct population require a complete re-training of the denoiser and generator network (B and C in Fig. 1).

### D. Experimental dataset

To test our procedure, we used the vim-1 dataset; a high quality, publicly available, recording of two subjects presented with natural images [1], [2]. The dataset, described in detail in [1], contains estimates of functional BOLD activity in response to greyscale natural photographs from voxels in visual brain areas V1, V2, V3, V4, V3A, V3B and LO and in voxels in visually responsive cortex anterior to LO. This dataset was acquired using a 4T INOVA MR scanner (Varian, Inc.) at a spatial resolution of 2mm × 2mm × 2.5mm and a temporal resolution of 1 Hz. During the acquisition, subjects viewed sequences of 20° × 20° greyscale natural photographs while fixating on a central white square. Photographs were presented for 1 s with a delay of 3 s between successive photographs. The data, available online at https://crcns.org/data-sets/vc/vim-1, is partitioned into distinct training and validation sets which consist of estimated voxel activation in response to 1750 and 120 photographs respectively. We further separated the validation set into a set of 100 samples used for inferring the noise parameters and a set of 20 samples for the final generative test. All figures in the current study use data from subject 1.

## Results and discussion

Before presenting our results for the encoding and generative procedure on the vim-1 dataset, we probe the effects of various observable properties of neural activity on the reconstruction accuracy on a synthetic neural dataset.

### A. Reconstruction variability depends on the encoding model accuracy, receptive field coverage and feature support

Our approach assumes that features reliably encoded in the brain-derived conditioning vector *C* will be present in the majority of samples from *p*(*X*|*C*), while features that are encoded noisily or not at all will vary across samples from *p*(*X*|*C*) just as much as they would in samples from *p*(*X*).

In order to validate this fundamental assumption, we first performed experiments with a set of synthetic brain activity measurements. The voxels in this synthetic dataset were randomly tuned to the feature maps of a DNN trained to discriminate digits in the MNIST digits dataset. We varied the noisiness of each synthetic voxel (i.e., the accuracy of its encoding model), the specific feature maps in the DNN to which the synthetic voxels were tuned (i.e., the voxels’ feature support), and the location and size of the synthetic voxels’ spatial RFs. We then examined the effects of these manipulations on the reconstructions generated by a GAN trained as specified in the methods section above. Specifically, we examined how the variability of samples from *p*(*X*|*C*) depends on the amount of noise, the feature support, and the RF coverage of the synthetic voxels.

The sample variability was estimated by calculating the (pixelwise or average) entropy from an estimate of the probability of a pixel being white or black across samples from 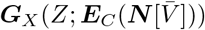. If all samples were identical, the entropy would be low, whereas highly variable samples result in high entropy. Note that low sample variability does not necessarily imply that the samples are accurate reconstructions (i.e. that they look like the image encoded in the conditioning vector *C*); nonetheless, we found that in these *in silico* experiments all reconstructions were clearly recognizable as the correct digit (unless otherwise noted).

Figure 2C-E illustrates the effects of manipulating the synthetic voxel populations on the reconstructed image samples. In all cases except where the feature support included only the first 2 convolutional layers (*V*_0_) of the DNN, the generator learned to produce convincing-looking digits of the category encoded. For the feature support *V*_0_, the reconstruction tended to be an amalgamation of edges that resembled digits but lacked overall coherence. On the other hand, these reconstructions were also less variable than those from feature support *V*_1_, *V*_2_ and *V*_3_, which suggest that the generator tuned specifically to noisy low-level visual features without consideration of the global arrangement.

We also tested the localization effect under the spatially homogeneous condition vector by restricting the RF support to only one visual hemifield. The result, shown in Fig. 2E, demonstrates a dependence on the local variability of the samples on the RF coverage of the encoded voxels. Furthermore, in this controlled experiment, the RF sizes relative to the visual feature of interest were extremely small. We thus also tested the effect of the size of the RFs by increasing all sizes by a factor of two and retrained the model. As expected, we observed that the variability of the samples increased with the RF sizes, which follows from the intuition that larger RFs cannot constrain the detail of the image as accurately.

These experiments with simple, synthetic brain activity confirmed that a conditional GAN could be trained to use stimulus information encoded in a brain-like code to narrow the variability of its samples around a target image. As expected, the variability in the samples was smallest for those features or those regions of the visual field most reliably encoded in the synthetic brain activities.

### B. Validation of the encoding model used to train the denoiser and the generator

As detailed above, we trained the denoiser and the ebGAN using surrogate data samples obtained from a voxel-wise encoding model. For the reconstruction to succeed at test time, when a real brain activity pattern is used, it is essential that the encoding model be accurate. Figure 3 shows that the encoding model used in this study accurately predicted brain activity measurements in response to natural scenes, and recovers well-known properties of receptive fields [13] and tuning along the visual hierarchy. Due to the low resolution of the input, the encoding model used here included a less rich set of features than in previous works [8]. This choice was made to accommodate the low-resolution images of the cifar-10 dataset used to train the denoiser and generator. In general—dataset permitting—we expect that one could use a high-resolution input for the encoding model while using down-sampled images for the generator in order to facilitate training of the latter. Nevertheless, the encoding model used here achieves comparable prediction accuracy in early visual areas and shows, as expected, the progressively increasing tuning to deep feature maps along the visual hierarchy also observed in Refs. [8], [14]–[16].

**Fig. 3.**
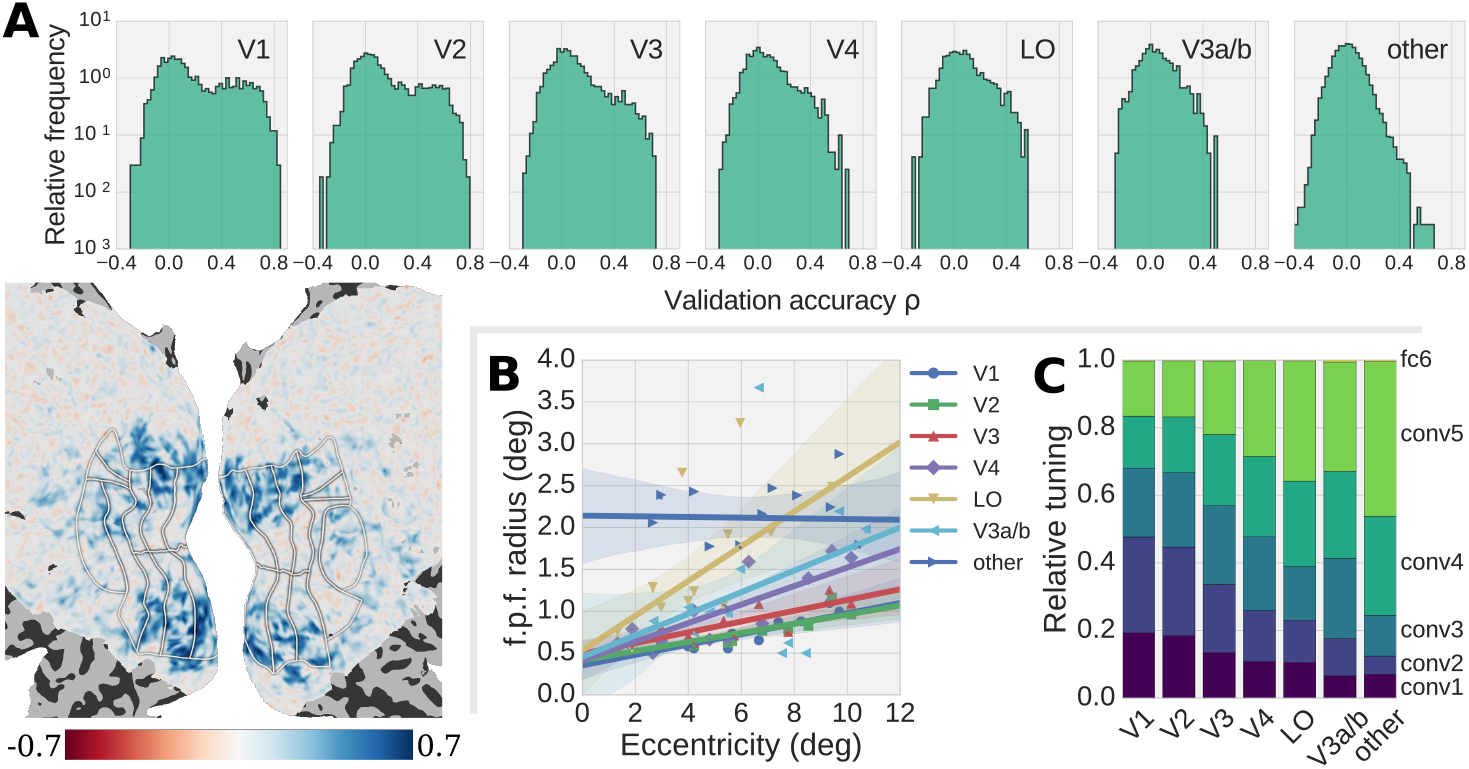
Encoding model. (A) shows histograms of model validation accuracy (i.e. the correlation coefficient between the prediction of the model and real voxel activity) for voxels in distinct visual areas (top right of each histogram). The inset displays model validation accuracy on a flattened map of visual cortex (subject 1 in the vim-1 dataset). The overall accuracy is lower than reported in [8]; however, the encoding model used here had many fewer model parameters, greatly decreased stimuli resolution, and was based on a DNN trained classify many fewer object categories. In spite of these suboptimal conditions, the model parameters still manage to reflect the dominant cortical organization, as seen in (B) and (C). (B) shows the estimated spatial RF size of voxels as a function as of its eccentricity (distance to fovea). (C) show the relative network layer contribution to the overall model accuracy for different visual areas. The network hierarchy aligns with the hierarchy of areas in the visual cortex.

Only voxels whose validation accuracy *ρ*_val_ were above a threshold of 0.2 were included in *V*. This corresponded to roughly 2900 voxels, 56% of which were found in early visual area V1-3, 21% in area V4, LO and V3a/b, and the remaining in visual cortex area anterior to LO.

### C. Reconstruction quality depends critically on voxel denoising

Before considering the fully cross-validated reconstructed images, we first examined reconstructions using the same set of brain activity patterns, *V*_val_, that were used to estimate the parameters of the noise model *N*. These brain activity patterns were used (indirectly) to construct the surrogate activity patterns on which the denoising autoencoder was trained. We found that these brain activity patterns are almost perfectly denoised by the autoencoder and therefore provide an upper bound on the reconstruction accuracy that we might achieve. Figure 4 shows a sample of stimuli reconstruction from brain activity measurements in *V*_val_. These demonstrate quite convincingly that the generator trained under the procedure described here learns to associate elements of the voxel predictions to low- and intermediate-level visual features to reconstruct the seen images. This suggests that extremely accurate reconstructions could be obtained with the proposed decoding strategy by adopting an experimental design that includes a set of brain activity measurements for the dedicated purpose of estimating noise parameters.

**Fig. 4.**
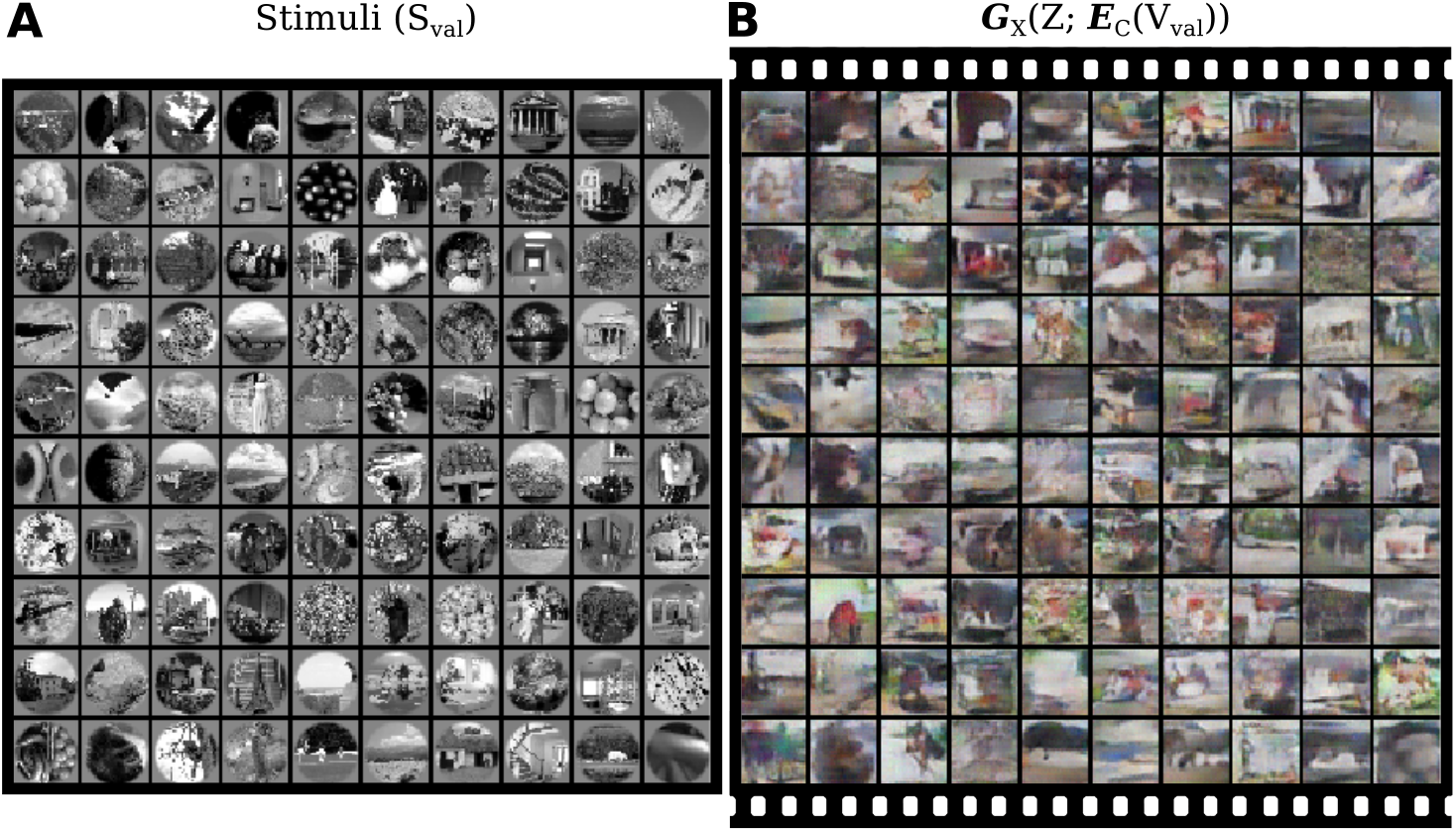
Image decoding from optimally denoised voxels. (A) The stimuli shown to the subject. (B) One sample from the generator conditioned on the evoked brain activity of each of the stimulus presentations. The colorization of the reconstruction samples are a result of the generator training dataset. For these reconstructions we used real measured brain activity patterns, but purposely used overfit parameters for the noise model used to train the denoising autoencoder. Thus, these reconstructions represent an upper bound on image reconstruction quality achievable with the proposed system, and suggests that proper estimation of noise model parameters is a critical step in our proposed decoding strategy. A composite video of samples demonstrating the stability of certain image elements is available online at https://github.com/styvesg/gan-decoding-supplementary.

### D. Reconstruction of visual stimuli from evoked brain activity

Figure 5 shows a few samples generated from conditioning on real held-out voxel activity measurements with the corresponding stimuli that was presented to the subject. While it would be difficult to identify the image from *a priori* considerations, one can observe that the dominant lines (horizon, mountains, etc.) appear to be preserved and that regions containing high spatial frequency details appear generally as such. A composite video of samples produces the impression of high variability of certain elements, while other, more robust, stand out as the main objects in the reconstructions.

**Fig. 5.**
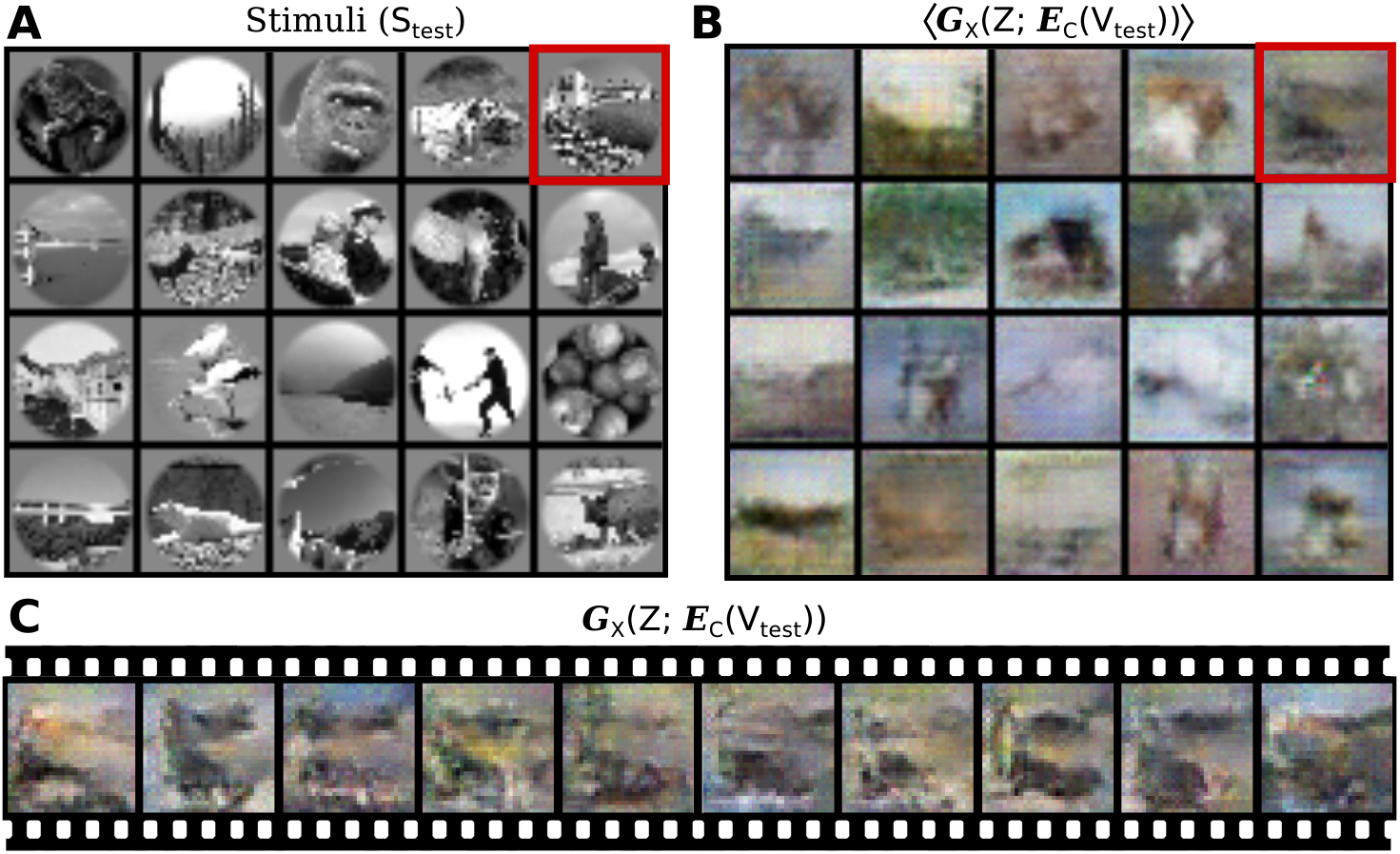
Generated samples. (A) Stimuli from the held-out test set *S*_test_. (B) The average over the reconstructed images recovers a coarse outline of the encoded stimuli. (C) 10 generated samples for the top-left stimuli (red square). It is also remarkable that the robust sample features are also preserved across re-training of all parts of the model. See https://github.com/styvesg/gan-decoding-supplementary for a composite video of such samples.

### E. Relation to other decoding methods

One way to model *p*(*X*|*V*) is to apply Bayes’ theorem with the encoding model describing the likelihood *p*(*V*|*X*) [2], [17]. This requires that one specifies an appropriate prior distribution *p*(*X*), which generally requires some fairly strong assumptions about *X*. For example, Refs. [2] and [3] used a large collection of natural images (videos) as a prior. The implicit representation of *p*(*X*|*V*) afforded by the GAN strategy does not require strong assumptions on the prior or posterior distributions, and thus greatly extends its expressiveness. Our results can be compared directly to those of Ref. [2] since we used the same experimental dataset. Averaged samples on the held-out set (Fig. 5) are comparable to the reconstructions presented in Ref. [2]; however, our method has the advantage of being able to produce multiple samples from *p*(*X*|*V*), as opposed to just the mean of the posterior. As we discuss below, this innovation helps to distinguish signal from noise in the reconstructed images (see supplementary videos for examples). Furthermore, the reconstructions in Fig. 4, where we assume near-optimal denoising, suggest that the ceiling on reconstruction quality might be much higher for the GAN decoding strategy presented here than for previous strategies.

Other methods center around a maximum likelihood estimation for the decoded image. One such obvious decoding procedure is to induce a generative process on *E_V_* (*X*) by performing gradient descent with respect to *X* under an objective that minimizes some distance *L*_rec_ between the neural predictions and measurements. However, such a process has several limitations. Mainly, the difficulty is that, while the pixel gradient ∇*_X_ L*_rec_ succeeds in reconstructing accurate images from low-level feature maps, it tends to drive reconstructions of deep feature maps into unnatural-looking images. This is related to the problem of interpretation of deep feature maps [18], [19] where several attempts have been made at fine-tuning prior-like constraints to the objective in order to favour certain aspects of the reconstructions, with limited success.

Shen *et al.* [20] attempted to improve the quality of the images obtained through such gradient-based method by adding a pre-trained generative model of images *E_V_* (*G_X_* (*Z*)) and perform gradient descent with respect to *Z* instead of *X* directly. This addition was intended to serve as a regularizer (or prior) to keep the generated image near the generative model manifold (of natural images). However, this procedure still does not permit an interpretation of the variance of the samples, even in principle. In contrast, instead of using an encoding model to score and tune the image generated *under* a pre-trained generative model, we trained a conditional generative model that learn the posterior distribution from brain to image *under* an encoding model.

### F. Toward generic decoding of mental images

It is important to conceptually separate the technical difficulties associated with the practical problem of image reconstruction from evoked brain activity from the theoretical problem of understanding the representative limitations of the processes that evoked this brain activity. The technical difficulties center around producing an inference procedure that can put together disparate and often corrupted pieces of information into a consistent whole. What we call consistent here refers to our prior knowledge of natural images.

Our analysis offers certain predictable prospects for decoding mental imagery (MI). First, it has long been known that evoked brain activity during MI does not cover the whole visual cortex uniformly, but tends to re-activate regions high in the visual hierarchy [21]. This implies, since these regions are best predicted by increasingly invariant features (Fig. 3), that an optimal generator would necessarily produce samples with high variability, as demonstrated by Fig. 2B. A recent fMRI study of MI revealed that population RF of voxels during MI tend to be larger in early visual area than during perception [22]. According to our results, this reduction in spatial specificity would also increase the variability of the generated samples, and suggests not merely a lower practical bound to MI reconstructions, but a lower theoretical bound as well.

## Conclusion

We have demonstrated the feasibility of using the GAN strategy to obtain an implicit representation of the probability distribution of images *p*(*X*|*V*) given observed voxels *V*. Our demonstration has focused on three elements: First, that common observable properties that are associated with neural activity (prediction accuracy, receptive fields and feature tuning) play a direct and obvious role in our capacity to reconstruct encoded stimuli under our method. Second, we maintain that the central challenge with our method is the voxel denoising strategy. Here we have used a model of the noise inferred from measurements and shown that a denoising autoencoder trained with surrogate data under this model effectively learns to denoise the voxel measurements. However, the relatively small size of the dataset meant that the noise model overfitted the noise, and thus generalized relatively poorly to held-out data. Nevertheless, the dominant features of the stimuli appeared to be recovered. Finally, we have shown that a generator conditioned on a brain-like code could be trained using purely synthetic data produced by a sufficiently accurate encoding model. This development permits extension of the generator to arbitrarily large natural image sets and clearly disentangles the problem of encoding, denoising and decoding.

## Acknowledgment

A preliminary survey of this work appeared in Ref. [23]. The authors would like to thank Jesse Breedlove for useful help and comments on the manuscript. This work was supported by grant NIH R01 EY023384 to TN.

